# Neutrophil arrest in myocardial capillaries drives hypoxia and impairs diastolic function in a mouse model of heart failure with preserved ejection fraction

**DOI:** 10.1101/2025.02.17.638717

**Authors:** Anne E. Buglione, David M. Small, Nathaniel H. Allan-Rahill, Tyler Locke, Laila M. Abd Elmagid, Rachel Kim, Soseh Hovasapian, Adina A. Mistry, Sofia A. Vaquerano, Chris B. Schaffer, Nozomi Nishimura

**Author notes:** co-first authors (Anne E. Buglione, David M. Small).

## Abstract

Impairments in myocardial blood flow have been recognized for decades in human patients with and animal models of heart failure with preserved ejection fraction (HFpEF), but the underlying mechanisms and roles in pathogenesis remain poorly understood. Using intravital cardiac microcopy in a ‘two-hit’ mouse model of HFpEF that combines high fat diet and inhibition of nitric oxide synthase, we identified an increase in slow or non-flowing neutrophils in capillaries compared to Control mice. In other mouse models of disease (e.g., in the brain of Alzheimer’s Disease mice), the presence of such ‘stalled’ neutrophils leads to organ-wide decreases in perfusion and oxygenation. Administration of antibodies against the neutrophil surface protein Ly6G to deplete neutrophils reduced the number of arrested neutrophils in myocardial capillaries, leading to improvements in myocardial hypoxia, diastolic function, and exercise capacity. This study identifies a previously uncharacterized cellular mechanism that explains myocardial blood flow deficits in mouse models of HFpEF, and demonstrates that improving myocardial blood flow improves cardiac function, without necessitating reversal of pathologic remodeling. Restoring myocardial perfusion by decreasing neutrophil arrest in myocardial capillaries may provide a strategy for improving heart function in HFpEF patients in the future.

## INTRODUCTION

Heart failure with preserved ejection fraction (HFpEF) is a category of heart failure characterized by impaired cardiac relaxation (diastolic dysfunction).^2^ Diastolic dysfunction leads to decreased ventricular filling and, consequently, decreased cardiac output. Patients present with a complex interplay of factors including reduced cardiac reserve, endothelial dysfunction, ventricular stiffening, and hypertension. The majority of therapies that have improved morbidity and mortality in heart failure with reduced ejection fraction (HFrEF) have proven ineffective in HFpEF.^3,4^ Treatment with inhibitors of the sodium-glucose cotransporter type 2 have been shown to reduce the risk of hospitalization in HFpEF patients. However, mortality.^5^ Further, the mechanism underlying this protective effect remains a mystery. Development of novel, rationally targeted therapeutic approaches in HFpEF mandates the development of a more complete understanding of alterations in cellular homeostasis occurring in this condition.

Multiple studies show that blood flow is decreased in myocardial tissue in human patients with HFpEF even in the absence of major arterial obstruction.^6-10^ The cause of this myocardial perfusion deficit is still unknown. However, evidence points toward decreased flow at a microvascular level as a potential cause. Reduced coronary flow reserve, defined as the ratio of coronary flow before and after stimulation, is common in HFpEF patients.^6-8^ Additionally, studies in HFpEF patients without obstructive coronary artery disease demonstrate increased microvascular resistance under stimulation,^9^ decreased volumetric flow,^6^ and reduced myocardial oxygen supply with exercise compared to healthy controls.^10^ Structural loss of capillaries via rarefaction is noted in patients^11^ and in animal models^12^ of HFpEF, and is thought to be related to inflammation.^13^ The microvasculature is often referred to as the likely origin of decreased flow, although the cellular cause and significance are unclear.

HFpEF in humans occurs due to low grade, chronic, systemic inflammation, often in the context of metabolic risk factors.^14,15^ It is not clear how inflammation in the periphery affects the heart and causes HFpEF, but multiple lines of evidence suggest a contribution of dysfunction of the coronary microvasculature.^16^ Endothelial activation is increased in HFpEF, indicated by the expression of vascular adhesion molecules and decreased bioavailability of nitric oxide (NO).^9,17-19^ Evidence supports immune and inflammatory cell infiltration of the heart, including neutrophils, in human and animal HFpEF^20^ and HFrEF studies^21^ of post-mortem tissue. However, the behavior of these cells within the cardiac microcirculation and how their recruitment impacts coronary microcirculatory dynamics have not been defined.

The convergence of the coronary microcirculation, inflammatory cells, and cardiomyocyte function in HFpEF phenotype requires *in vivo* study. Intravital 2-photon microscopy is uniquely able to show the dynamic interactions of circulating leukocytes within the microvasculature in living animals.^22-26^ Recent technical advances have overcome the challenges associated with imaging the live beating heart in mouse models of disease.^1,27-29,30^ This enables direct visualization of dynamic processes such as blood flow, which are impossible to study in extracted tissues, cultured cells, or external models such as the Langendorff perfusion. These imaging capabilities provide a link between microvascular dysfunction, inflammation, and reduced blood flow in the HFpEF heart. Using intravital cardiac two-photon microscopy in mouse models of heart failure, we show that neutrophils arrest in coronary capillaries to reduce microvascular blood flow, contributing to cardiac remodeling and dysfunction. The elimination of these stalled capillaries is associated with rapid improvement of hypoxia and diastolic dysfunction, without necessitating reversal pathologic remodeling.

## RESULTS

### Two-photon intravital imaging shows an increase in myocardial capillaries with slow moving luminal neutrophils in HfpEF

HFpEF was induced in male and female C57Bl/6 mice using a combination of high-fat diet and L-NAME in drinking water for 15 weeks (‘HFpEF’) and compared to age and sex matched controls who remained on a standard chow diet and water (Control) (Fig. 1a). HFpEF mice developed ventricular hypertrophy, which was readily apparent both grossly and assessed by the ratio of heart weight to tibia length (Fig. 1b). Mitral E/E’, a measure of diastolic dysfunction, was increased in HFpEF mice, while left ventricular (LV) ejection fraction was unchanged, as previously reported (Fig. 1c)^11.^ To investigate the interplay between microvasculature, leukocyte activity, and myocardial capillary blood flow in HFpEF, we used intravital cardiac two-photon microscopy to image the left ventricular epicardial microvasculature in anesthetized, ventilated mice (Fig. 2). ^26,27,3^. Plasma was labeled with retro-orbital injection of dextran-conjugated dyes or quantum dots. Erythrocytes and leukocytes are visible as regions of excluded labeling, which appear as dark patches within the vessel. Use of Catchup^IVM-Red^ mice, which express TdTomato on the surface of neutrophils,^31^ and heterozygous knock-in mice (Cx3Cr1^GFP/+^ x CCR2^RFP/+^)^30,31^, which express green and red fluorescent proteins (GFP and RFP) in monocytes and macrophages, allowed identification of specific classes of leukocytes (Fig. 3a and b). Labeled neutrophils and monocytes within capillaries were manually tracked in image stacks (100 frames at 30 fps from approximately 20 to 200 µm below the left ventricular surface in 2 µm steps). Due to the periodic motion from heartbeat and breathing at approximately 5-6 Hz and 1.6 Hz, capillaries appear repeatedly in the imaging stack and are recognizable over about 1-2 minutes, allowing visualization of neutrophils and monocytes as they transit through the capillary beds. While the majority of neutrophils and monocytes were visible for less than ∼2 s, many cells took substantially longer to traverse the 340x 340 µm imaging region. These stalled neutrophils and monocytes appeared in the same location over multiple image frames (Fig. 3a and b), sometimes apparently not moving at all over several minutes of imaging.

**Figure 1.**
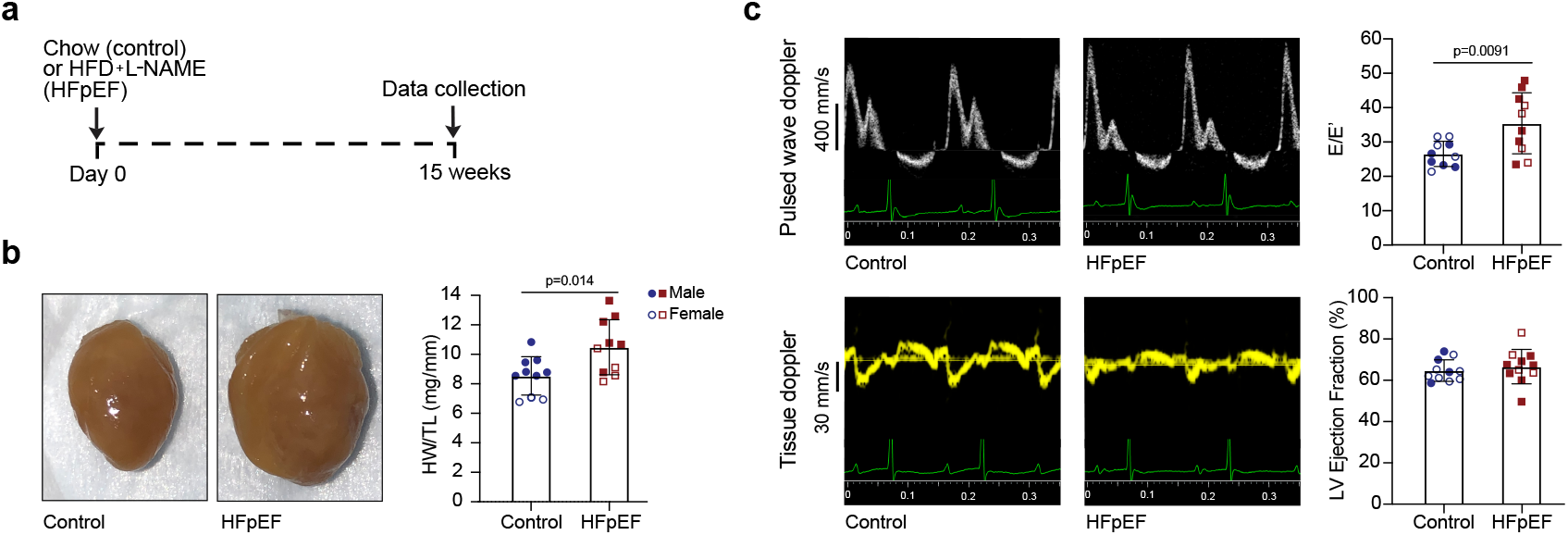
Mouse model of heart failure with preserved ejection fraction. (a) Schematic of HFpEF induction protocol (top). Age and sex matched animals received high fat diet (HFD) and L-NAME ad lib (HFpEF) or normal chow and drinking water (Control) for 15 weeks, beginning at age 10-12 weeks old. Example images (bottom left) of extracted hearts from control and HFpEF mice, showing gross cardiac hypertrophy. Plotted (bottom right) ratio of heart weight (HW) to tibial length (TL) ratios in control and HFpEF animals, demonstrates increased cardiac mass, controlling for body size. (b)Representative pulsed-wave Doppler (top left) and tissue Doppler (bottom left) traces taken at the level of the mitral valve. Plotted ratio of mitral E wave to mitral E’ wave (E/E’ ratio, top right). Left ventricular ejection fraction (bottom right). Data points represent individual animals and include wildtype mice or mice with labeled neutrophils or monocytes.

**Figure 2.**
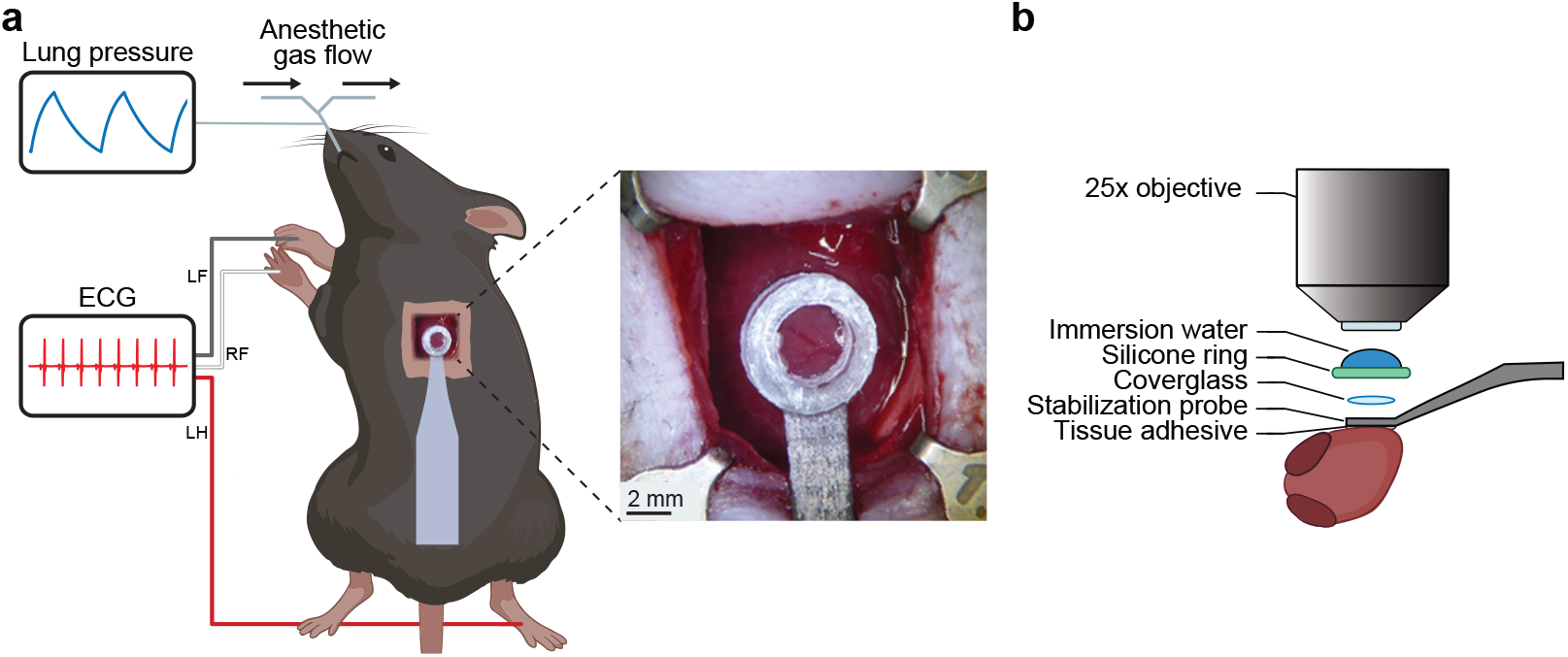
Surgical and imaging approach to the intravital two-photon microscopy in the beating mouse heart. (a). The left ventricle is accessed by performing a thoracotomy on an intubated, mechanically ventilated mouse. A three-lead (LF: left forelimb; RF: right forelimb, LH: left hindlimb) electrocardiogram (ECG) records the electrical activity in the heart. ECG and lung pressure are monitored and recorded throughout surgery and imaging. Inset: the heart surface showing coronary surface vessels. (b). A 3-mm glass window is stabilized by a titanium probe and attached to the heart surface with tissue adhesive^1^.

**Figure 3.**
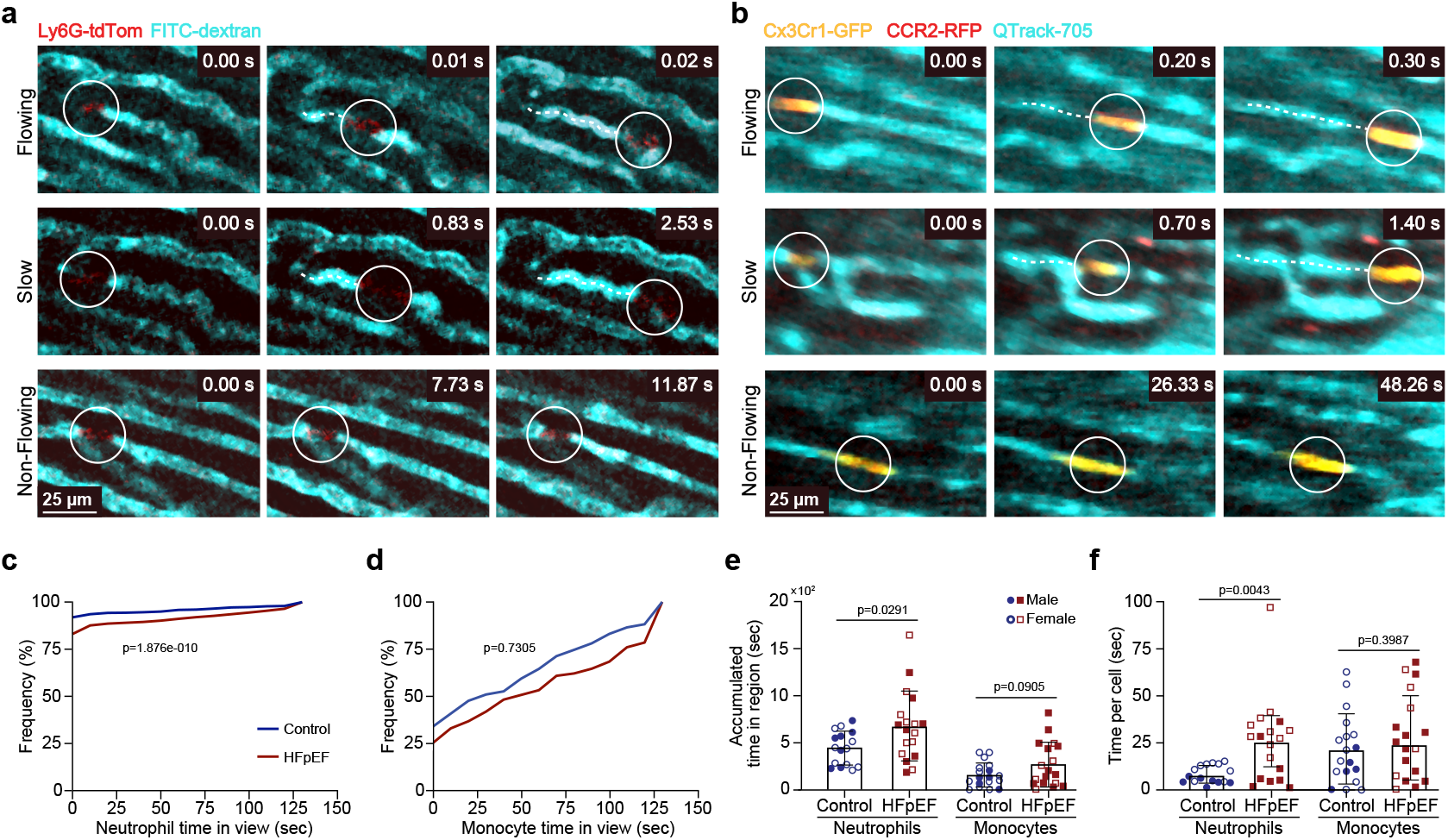
Mouse model of HFpEF leads to increased neutrophil residence in myocardial capillaries compared to Control animals in C57Bl/6 mice. (a) Representative in vivo, two-photon (2P) microscopy images of myocardial capillaries and capillary neutrophils in the left ventricular wall in a HFpEF mouse expressing tdTomato (red) with the Ly6G promoter. Blood plasma is labeled with FITC-dextran (cyan). Scale bar = 25 µm. (b) Representative in vivo 2P microscopy images in a HFpEF mouse expressing red fluorescent protein (RFP, red) driven by the CCR2 promoter and green fluorescent protein (GFP, yellow) with the Cx3Cr1 promoter in monocytes. Plasma labeled with quantum dots (Qtrack-755, cyan). Scale bar = 25 µm. (c) Distributions of time for which Ly6G -tdTomato neutrophils were present in the visible region of image stacks in Control and HFpEF mice. p=1.87 x 10^-10 (d) Distributions of time for which Cx3Cr1-GFP monocytes were present in the visible region of image stacks in Control and HFpEF mice. p=0.735 (e) Accumulated time for which Ly6G -tdTomato neutrophils or Cx3Cr1-GFP monocytes were present in the visible region of image stacks, summed over all cells in each stack. Each data point represents an individual stack (f) Average time for which individual Ly6G -tdTomato neutrophils or Cx3Cr1-GFP monocytes were present in the visible region of image stacks. Data points represent the mean time for all cells in each stack. Control/neutrophils n = 6 mice, HFpEF/neutrophils n = 4, Control/monocytes n = 4, HFpEF/monocytes n = 6. Each data point in (e-g) represents an image stack. Indicated p-values are for Student’s t-test unless otherwise noted

In Catchup mice, we measured the time for which each neutrophil was visible within the image stack. To reduce the variability due to the differing amount of time a particular capillary is visible in the image volume, we sampled over multiple image stacks per mouse. The distribution of times differed between HFpEF and Control, with a greater proportion of neutrophils in HFpEF persisting for longer times (p<0.0001 Kolmogorov–Smirnov (KS) test, Fig. 3c). The total amount of time that one or more neutrophils was present in the image stack (accumulated time in region) and the time visible per cell were increased by 1.5x and 3.1x (Fig. 3e and 3f) in HFpEF relative to Control mice. Similar experiments using Cx3Cr1^GFP/+^ x CCR2^RFP/+^ mice with labeled monocytes^23,30,32^ revealed no difference between the monocytes of HFpEF and Control mice in the distribution of time in the image stack (p= 0.73 KS test, Fig. 3d). The majority of cells visualized expressed both Cx3Cr1-GFP and CCR2-RFP (Fig. 3b). Cells expressing GFP were tracked as a measure of monocytes. Monocytes were in view for longer than were neutrophils, regardless of treatment group (Figure 3e and f). However, far fewer monocytes (n= 148 cells across 9 Control mice, n= 198 cells across 9 HFpEF mice) were identified than neutrophils (n= 1184 cells across 8 Control mice, n=1028 across 8 HFpEF), regardless of treatment group. Although the average time for which individual monocytes were visible was greater than the average time for which individual neutrophils were visible (Fig. 3f), the difference in duration between Control and HFpEF mice was not statistically significant. Further, monocyte stalls occurred less frequently than neutrophil stalls, resulting in an overall lower accumulated time for monocytes (Fig. 3e).

Subsequent experiments focused on neutrophils because of the clear effect of treatment group on the distribution of time for which cells were present in view (Fig. 3c), and the larger accumulated time in capillaries compared to monocytes (Fig. 3e). The number of neutrophils visible for ≥ 100 seconds, or ‘stalled,’ was doubled in HFpEF versus Control (Fig. 4a). Correspondingly, the incidence of ‘flowing’ neutrophils, defined as having a time in view of < 5s, was lower in HFpEF mice compared to Controls. HFpEF transit times were shifted towards longer time in view (Fig. 4b and c), reflecting slower overall movement. Although there are reports of loss of capillaries in HFpEF, we found total capillary length, measured from z-projections spanning 20-µm of the imaging volumes, to be similar in HFpEF and Control mice (Fig 4d and e). Total number of neutrophils per minute normalized by capillary lengths was also similar in HFpEF and Control (Fig. 4f). The total time for which neutrophils were visible was greater in HFpEF than Control and there were fewer cells per minute, suggesting that capillaries of HFpEF mice are obstructed by neutrophils for longer durations compared to capillaries of Control mice. On the whole, neutrophils were stalled and slowed in capillaries more in HFpEF than in Control mice. Only one interstitial neutrophil was identified during in vivo imaging out of all neutrophils counted (1208 in HFpEF, 1184 Control). To further localize interstitial neutrophils, we used immunolabeling against tdTomato and quantified neutrophils in ventricular tissue sections. Cryoinfarction was used as a positive control. While there was a trend towards an increase in the number of total and interstitial neutrophils HFpEF mice, this elevation is negligible compared to the number of interstitial neutrophils adjacent to an infarction^29,33-36^ (Fig. 4g-i).

**Figure 4.**
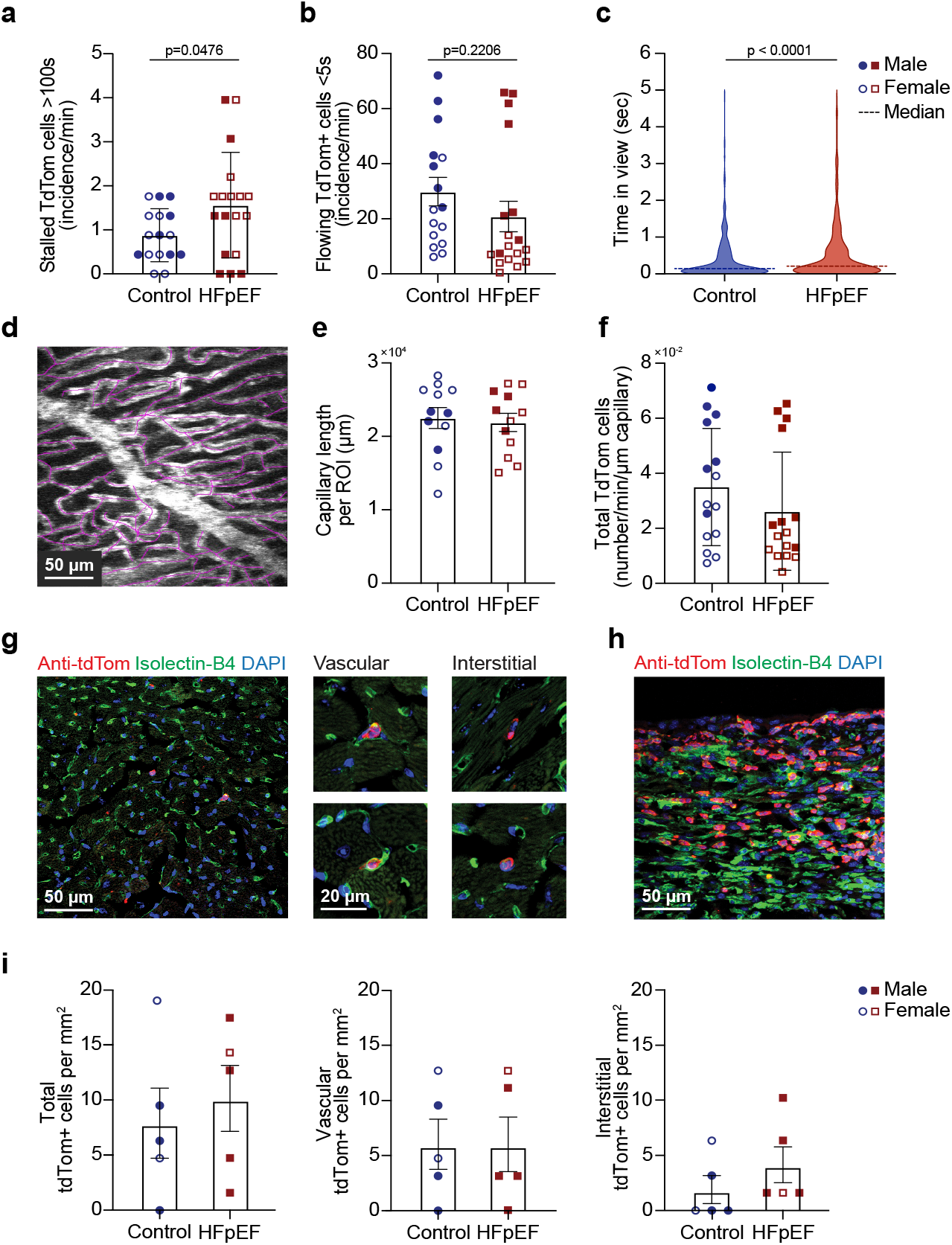
Characterization of myocardial capillary neutrophils in Catchup mice. (a) Number of neutrophils per time visible in two-photon microscopy image stacks for 100 seconds or longer (categorized as ‘stalled’) or (b) less than 5 seconds (categorized as ‘flowing’). (c) Distributions of time in view for ‘flowing’ neutrophils. Dashed lines represent the median. (d) Capillaries were traced in 20 µm projections of images from two photon image stacks to calculate (e) summed linear length of capillaries (f) Incidence of neutrophils observed in an image stack per minute normalized by total capillary length (g) Example images of confocal microscopy images of sectioned left ventricular tissue taken from a HFpEF phenotype mouse. Neutrophils were detected by an antibody against tdTomato (anti-tdTomato, red). Vessel wall is labeled with isolectin-B4 (green) and nuclei with DAPI (blue). Neutrophils were identified by co-localization of anti-tdTom and DAPI and were categorized as vascular when they were adjacent to isolectinB4-labeled vasculature and as interstitial when no contact was observed. (h) Left ventricular wall tissue collected 7-days after a cyroinfarction. This served as a positive control for histological neutrophil labeling. Labeling is the same as (g). (i) Quantification of all visible (left), vascular (middle), and interstitial (right) neutrophils in tissue sections from Control and HFpEF mice. Labeling is the same as (g).

### Hypoxia, diastolic function, and fibrosis are rescued by depleting neutrophils for 4 weeks in late stages of HfpEF

We tested the effects on disease progression of long-term stall removal by depleting neutrophils for 4 weeks, beginning at week 11 of HFpEF induction. An antibody against the neutrophil surface protein Ly6G (α-Ly6G) was administered intraperitoneally at a dose of 2mg/kg every 3 days for 4 weeks as previously published to maintain neutrophil depletion^37^. An isotype control (denoted ‘Iso’ in Figure 5) was administered with the same timing and dosage. α-Ly6G treatment did not affect weight gain (Fig. 5b), cardiac hypertrophy (Fig. 5c) or left ventricular ejection fraction (Fig. 5d) compared to isotype control. However, in HFpEF mice, treatment with α-Ly6G led to recovery of diastolic function (Fig. 5e) with E/E’ similar to that of Control. Similarly, myocardial hypoxia, measured by the pimonidazole immunoassay, increased in HFpEF hearts to 150% of Control mice (Supplementary Figure 4). 4-week treatment of HFpEF mice with α-Ly6G reduced hypoxia to nearly the levels of Control mice (Fig. 5f and g). E/E’ and pimonidazole fluorescence were not affected by α-Ly6G treatment in Control animals when compared to Control animals treated with isotype control antibody (Fig. 5e-g). Collagen, as assessed by Masson’s trichrome, was less increased in HFpEF animals treated with α-Ly6G than with isotype control compared to Control animals treated with either α-Ly6G or isotype control (Fig. 5h-i).

**Figure 5.**
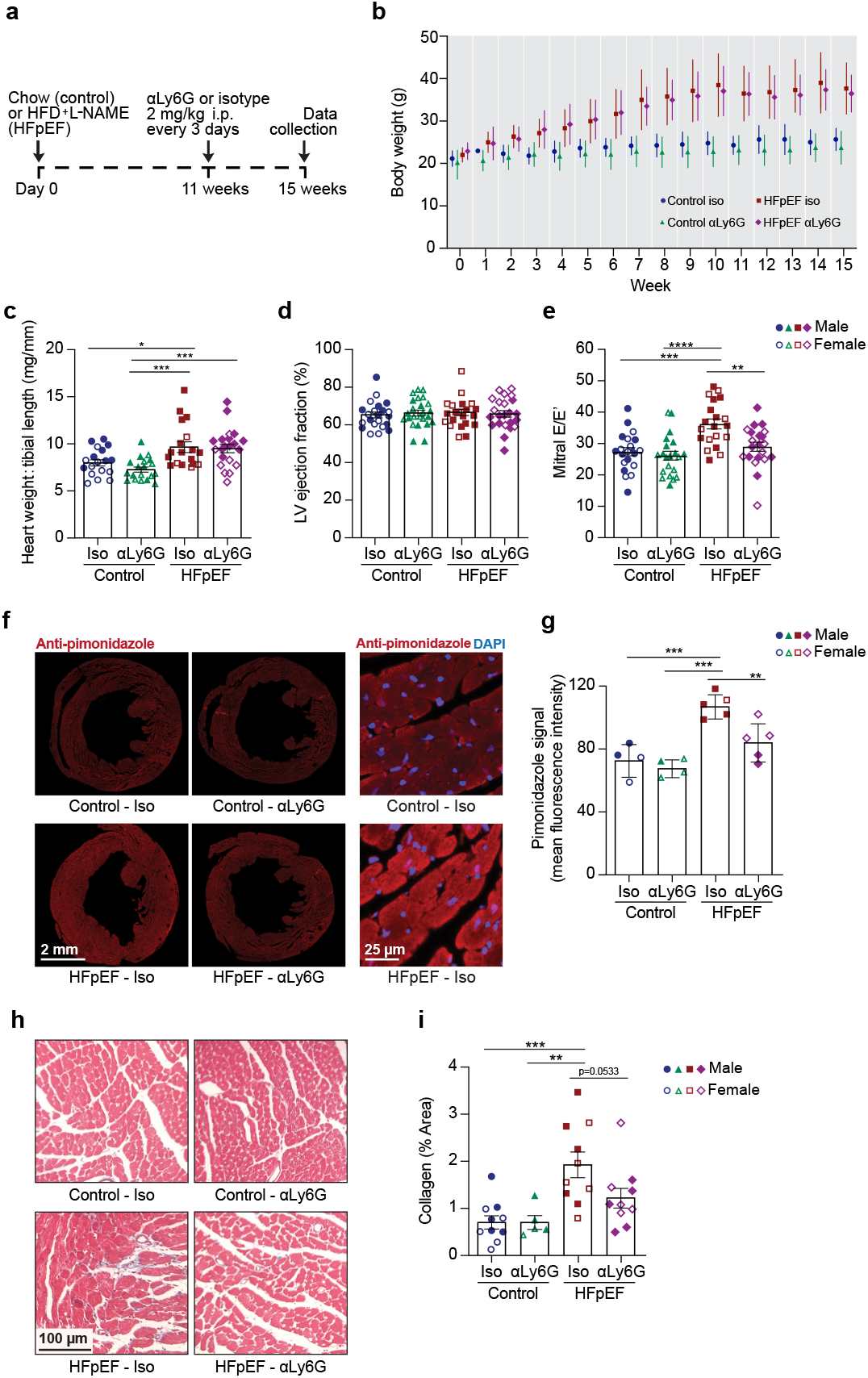
Long-term neutrophil depletion improves diastolic function and fibrosis, without affecting weight gain or cardiac hypertrophy. (a) Schematic of HFpEF induction and antibody treatment protocol. Age and sex matched animals received high fat diet (HFD) and L-NAME *ad lib* (HFpEF) or normal chow and drinking water (Control) for 15 weeks, beginning at age 10-12 weeks old. Starting at week 11 of the experimental protocol, mice were injected with antibodies against Ly6G (a-Ly6G) or a nonspecific isotype control antibody (Iso) every three days until imaging and tissue collection. (b) Animal weights starting from initiation of HFpEF or Control protocols to Week 15. Antibody treatments were started at week 11. (c) Heart weight normalized to tibial length in HFpEF and Control mice treated with a-Ly6G or Iso antibodies (d) Left ventricular (LV) ejection fraction in HFpEF and Control mice treated with a-Ly6G or Iso antibodies (e) Mitral E/E’ ratio in HFpEF and Control mice treated with a-Ly6G or Iso antibodies. (f) Example fluorescence microscopy of ventricular section from animals injected with a marker of hypoxia (pimonidazole), detected with anti-pimonidazole (red) (g) quantification of anti-pimonidazole labeling (h) Representative images showing Masson’s trichrome staining of collagen in left ventricular tissue sections (i) Quantification of Masson’s trichrome staining All data collection took place at week 15 of the treatment period. Student’s t-test, * p<0.05, ** p<0.005, ***p<0.00050

### Acute neutrophil depletion in late-stage HFpEF rescues hypoxia, diastolic function, and reduces stalled capillaries

We tested the effects of acute (24 hour) depletion at an advanced stage (week 15) of HFpEF (Fig. 6a). Using flow cytometry, we found that αLy6G (4 mg/kg IP once) led to a 54% reduction in systemic circulating neutrophils by 24h compared to mice administered an isotype control antibody (Fig. 6b and c). We took advantage of this rapidity to isolate the effects of neutrophil depletion from possible slower changes at the end of the 15-week HFpEF induction (or Control) in Catchup mice. Repeated echocardiography before and 1 day after αLy6G treatment revealed that treatment reduced E/E’ in HFpEF to the same value as in Control mice, while both HFpEF mice receiving isotype and Control mice receiving αLy6G did not change (Fig. 6d). Myocardial hypoxia, measured by the pimonidazole immunoassay, increased by 1.5-fold in HFpEF hearts compared to Control (Fig. 6e-f). One day following αLy6G administration, pimonidazole labeling decreased in HFpEF mice compared to isotype control treatment, approaching the levels found in Control. Intravital cardiac two-photon microscopy in Catchup mice with 15-week HFpEF induction showed that 1 day after αLy6G, the rate of neutrophils appearing in image stacks was reduced to less than half the rate with the isotype antibody (Fig. 6h). This change was statistically significant despite apparent inadequate neutrophil depletion in one mouse (two stacks, denoted with asterisks in Fig. 6h).

**Figure 6.**
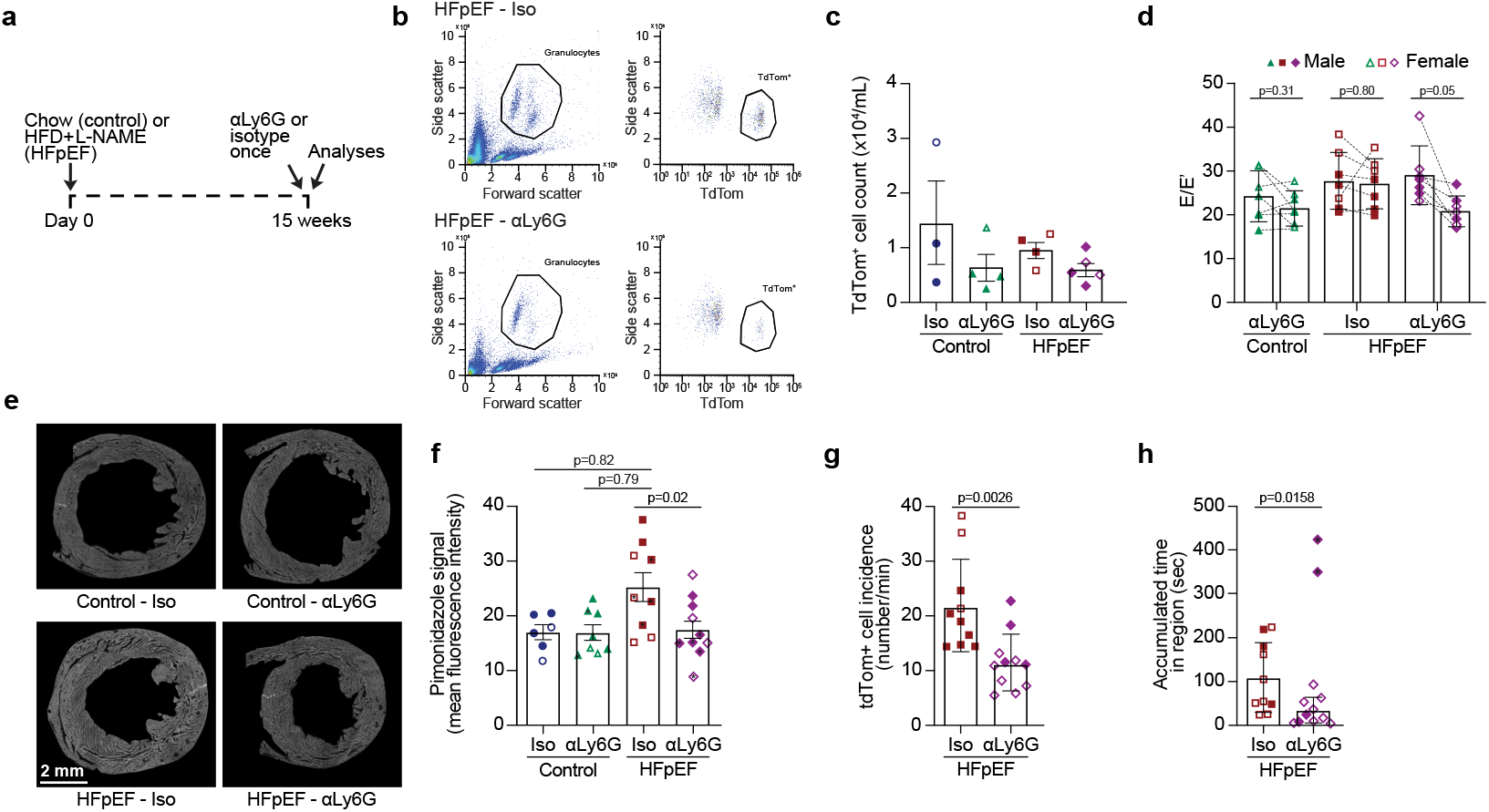
Acute, 1-day neutrophil depletion improves diastolic function and hypoxia and reduces neutrophil obstruction in capillaries. (a) Animals received high fat diet and L-NAME for HFpEF induction (HFpEF), while control animals received normal chow and water for 15 weeks. One day before imaging and tissue collection, animals were injected with antibodies against Ly6G (aLy6G) or an nonspecific isotype matched antibody as control (Iso). (n = 4 per group) (b) Representative flow cytometry data for blood drawn from HFpEF Catchup mice 24 hours after treatment with isotype control antibodies (top row) and α-Ly6G (bottom row). The left column shows forward and side scattering from the entire population of blood cells (after lysis and removal of red blood cells). The circled region indicates granulocytes, including neutrophils. The second column shows the gate on TdTom+ cells, indicating neutrophils (c) quantification of circulating neutrophils in Catchup HFpEF mice after 1day of treatment with anti-Ly6G or isotype control antibodies. (d) Echocardiography performed before and 1 day after anti-Ly6G or isotype control in HFpEF and healthy mice to measure diastolic function with E/e’. Paired t-tests, control-aLy6G: n = 6, HFpEF-Iso: n = 6, HFpEF-aLy6G: n = 7 (e) Example confocal microscopy of sections from control and HFpEF mice 1 day after a single dose of with either isotype control or anti-Ly6G antibodies. Pimonidazole, a marker of hypoxia was injected 90 minutes before euthanasia, and identified using immunohistochemistry. Greater fluorescence indicates greater hypoxia (f) Quantification of fluorescence using antibodies against pimonidazole (g) In vivo two-photon microscopy of Catchup mice was used to quantify the incidence of tdTomato-expressing (tdTom) neutrophils and (h) the sum of times a neutrophil was visible in image stacks 1 day after anti-Ly6G or isotype injection. (n = 5-6 mice). Each data point represents an image stack.

## DISCUSSION

### Capillary occlusions drive hypoxia in HFpEF

While many have speculated that microvascular dysfunction contributes to the heart failure, the mechanism has been elusive in part because it was difficult to visualize. Recently developed intravital two-photon microscopy enables imaging of blood flow in myocardial capillaries. In a mouse model of HFpEF, this imaging revealed that a subset of capillaries was transiently occluded primarily by arrested neutrophils that stall or slow blood flow (Fig. 3). This dynamic phenomenon was not reflected in histological counts of neutrophils in vessels and involves a tiny fraction of the number of neutrophils recruited to tissues after acute injuries such as infarctions (Fig. 4), explaining why it may not have been recognized previously without in vivo imaging. In HFpEF mice, the improvement of a marker of hypoxia just 24 hours after a single treatment of anti-Ly6G (Fig. 6) suggests that the presence of stalled neutrophils decreases overall myocardial oxygenation, and that this effect is reversible with removal of stalls.

### Diastolic function is rescued independently of remodeling and hypertrophy

Diastolic function also improved with the acute, 1-day treatment to reduce stalling of capillaries, providing benefits similar in magnitude to an extended 4-week treatment. Both long and acute treatment improved E/E’ to the levels of healthy, sham-treatment mice, but the 4-week treatment also had an additional benefit of reducing fibrosis, as measured by collagen staining. Although conventional wisdom suggests that remodeling and the accumulation of fibrosis drives the inability to relax by increasing tissue stiffness, the success of the acute treatment suggests a myocardial relaxation mechanism related to blood flow improvement. One possibility that relates diastolic stiffness to hypoxia on a fast time scale is the fact that low oxygenation can impair the conversion from ADP to ATP and high ADP levels can impair crossbridge detachment, prevent acti-myosin unbinding^38^. Reminiscent of the results here, we recently discovered using similar two-photon microscopy to this work, that a tiny fraction of capillaries (1-2%) in the brains of Alzheimer’s mouse models are stalled by arrested neutrophils, and this leads to a 25% decrement in brain blood flow^39^. This flow reduction was sufficient to contribute to cognitive dysfunction, demonstrated by the extremely rapid (3 hours) onset^39^ of short-term memory rescue caused by reducing the capillary stalls with α-Ly6G (same as used in this study). In both the Alzheimer’s and HFpEF models, long-term treatment even at late stages resulted in sustained functional improvement ^40,41^. The two parallel results suggests that the cumulative effects of distributed microscale occlusions can have large impacts on overall organ function and that at least some of effects can be rapidly reversed despite tissue remodeling.

### Evidence from patients and experimental work provide support for neutrophil adhesion in capillaries

Patient samples show elevated expression of the adhesion markers ICAM1, VCAM1, and E-selectin^17,18,42^, which could contribute to the arrest and slow flow of white blood cells in myocardial capillaries. HFpEF patient samples also show capillary rarefication^13^, suggesting endothelial cell damage that could drive white cell arrest^43^. Molecular methods implicate reactive oxygen species (ROS) and downstream signaling in cardiomyocytes and fibroblasts^16,44^. ROS also affect endothelial cells and drive increased inflammation as seen through increased expression of adhesion molecules^18,42^ and changes in barrier function^45^. In HFpEF patients, the neutrophil to lymphocyte ratio, a marker of inflammation, correlated with poor prognosis^46^. Myeloperoxidase (MPO) levels were increased in HFpEF patients, also suggesting an involvement of neutrophils^47^. Neutrophil depletion can have deleterious consequences after a myocardial infarction^48^, suggesting that after acute injury, neutrophil ‘clean up’ roles may be critical^49^. However, it has also been shown that depleting neutrophils in the pressure overload model of HFrEF can prevent heart failure^21^. The link between neutrophils and fibrosis is supported by additional work in mice. Zhang, et al. found similarly improved E/E’ and fibrosis without rescue of myocardial hypertrophy in a mouse model of HFpEF treated with the SGLT2 inhibitor drug dapagliflozin^50^. Similar changes occurred following in vivo degradation of neutrophil extracellular traps (NETs) with deoxyribonuclease 1. Further, in cultured neutrophils, dapagliflozin led to less NET release following stimulation with a NET-inducing compound.

### Future work

Follow up studies exploring the idea that perfusion deficits caused by capillary stalls contribute to pathologic changes in HFpEF are needed to address the limitations in this study. First, while α-Ly6G is useful as experimental tool, it is not expressed by human neutrophils. Further, long-term neutrophil depletion would be neither practical nor safe. While some studies have shown that α-Ly6G results in incomplete depletion of some subsets of neutrophils.^51^ However, the results presented here and data from the brains of mouse models of Alzheimer’s disease^39-41^ show significant changes in multiple physiological parameters, suggesting that the degree of depletion is biologically meaningful or at minimum warrants further work using different strategies. A similar phenomenon of stalled and slowed capillaries in an alternative model of heart failure ApoE^-/-^ with high fat diet, suggests that this phenomenon is not an artifact of the HFpEF model (Supplementary Figs. 2-3). Given that this study was not powered to resolve sex differences, the absence of notable sex dependence in stall incidence or functional recovery with depletion should be interpreted with caution. Further studies on sex difference, particularly incorporating menopause models, will be critical in assessing the translational importance of these results.

## CONCLUSION

This work points to stalled blood flow in myocardial capillaries due to transient arrest of neutrophils as novel mechanism underlying decreased myocardial blood flow in HFpEF. Antibody-mediated elimination of neutrophils and, consequently, obstructions resulted in less hypoxia, fibrosis, and diastolic dysfunction. The mechanism underlying slowed transit of neutrophils is not known. However, because neutrophil adherence to the endothelium occurs as the first step of neutrophil extravasation during inflammation, such capillary stalling with likely occurs in many conditions. Although antibody-mediated neutrophil depletion is not a practical or safe treatment option in humans, the process of neutrophil arrest may represent a new target for future therapies. Modulating stalling may also be an unknown mechanism underlying the benefit of existing treatments, such as SGLT2 inhibitors, for which the mechanism of protective effect is unknown.

## METHODS

### Mice

All animal procedures were approved by the Cornell Institutional Animal Care and Use Committee (protocol number: 2015-0029) and were performed under the guidance of the Cornell Center for Animal Resources and Education. All mice were on a C57BL/6J background. Catchup^IVM-red^ mice were generated by crossing homozygous Ly6G-Cre-tdTomato (Catchup)^34^ mice to a homozygous tdTomato reporter line from a Cre-activatable CAG promoter in the *ROSA26* locus (Ai9, Jackson Laboratory (Jax) stock #007909; ROSA-CAG-tdTom). These mice were crossed to wild type mice and offspring with one intact Ly6G allele were used for experiments. Cx3Cr1^+/GFP^-CCR2^+/RFP^ mice were generated by crossing homozygous B6.129P(Cg)-Ptprca-Cx3cr1tm1Litt/J (Jax #005582)^35^ with homozygous B6129(Cg)-Ccr2tm2.1lfc/J (Jax #017586)^31^ mice. Mice deficient in both copies of the ApoE allele (ApoE^-/-^) were purchased from Jackson Laboratory (Jax #002052)^52^ and bred locally.

### Heart failure models

All experiments used both male and female mice. Mice were maintained on a 14-h light (6:00 to 20:00), 10-h dark cycle with ad lib access to food and water. Animals in the Healthy Control group received standard diet (2916, Teklad) and plain drinking water. The HFpEF phenotype was induced starting at age 8-12 weeks for the indicated periods of time using high fat diet (D12492, Research Diet, 60% calories from lard) and *N*^ω^-nitro-arginine methyl ester (L-NAME, 0.5 g/L, Sigma Aldrich) in drinking water. The pH of the L-NAME solution was adjusted to 7.4 using sodium hydroxide. The L-NAME solution was stored in light protected bottles and prepared every 1-2 weeks. All ApoE^-/-^ mice used in experiments were fed a high fat diet (Harlan, TD88137) for 20 weeks starting at age 8-12 weeks.

### Surgical approach to the beating heart

Intravital imaging was performed in anesthetized, mechanically ventilated mice. Anesthesia was induced with buprenorphine HCl (0.1mg/kg, Par Pharmaceutical, in saline) subcutaneously, and ketamine/xylazine (50mg/kg, Ketaset; 5mg/kg, Covetrus, as a combined solution in saline) intraperitoneally. A 22-gauge, 1-inch polyethylene catheter was inserted to a depth of approximately 2cm past the maxillary incisors and connected via luer-lock to a ventilator (CWE Inc. SAR-830/P ventilator) to provide positive pressure ventilation (95 breath/min, 12-18 cm H_2_O end-inspiratory pressure, 2 cm H_2_O PEEP, inspiratory/expiratory ratio of 1:1). Anesthesia was maintained with 1-3% isoflurane in 100% oxygen. Mice were positioned in right lateral recumbency on a feedback-controlled heating pad (40-90-8D DC, FHC) to maintain body temperature at 37°C. Glycopyrrolate (0.5mg/kg, Baxter, Inc) was administered intramuscularly into the caudal thigh. Glucose (5% w/v in saline, 10mL/kg) was administered subcutaneously. Electrocardiogram (ECG) leads were connected subcutaneously in the distal forelimbs and left hindlimb and connected to an isolated differential amplifier (World Precision Instruments ISO-80). Hair on the ventral thorax was removed with Nair, and the surgical site cleaned with ethanol. A skin incision was made starting 1 cm lateral and cranial to the xiphoid process, and extended 1-cm laterally. Subcutaneous tissue was bluntly dissected to expose the pectoral muscles, which were transected and reflected to expose the ribs and intercostal muscles. Ribs 7 and 8 were separated with forceps and retracted with either metal retractors or stay sutures. Adipose tissue surrounding the heart was carefully removed to provide 4 mm of exposed surface area. The pericardium was lifted, incised, and removed to allow stable adhesion of the titanium window support^31^ to the heart surface. Tissue adhesive (3M, Vetbond) was applied to the bottom of the titanium window support, which was then lowered to the surface of the heart, as described previously. Throughout surgery and imaging, lung pressure and ECG were continuously monitored and recorded.

### Intravital cardiac multiphoton microscopy

A silicone O-ring (4mm outer diameter, 3mm inner diameter) was glued to the top of the titanium imaging support, surrounding the window. The ring was filled with either distilled water or refractive index matching gel to immerse the microscope objective (Olympus XLPlan N 25 x 1.05 NA). Imaging was performed using a custom-built raster-scanning microscope equipped with four detection channels and an 8-kHz resonant scanner (Cambridge Technology). Data was acquired using a National Instruments digitizer (NI-5734), FPGA (PXIe-7975), and multifunction I/O module (PXIe-6366) for device control, mounted in a PXI chassis (PXI-1073) controlled by ScanImage 2019b. A Ti:Sapphire laser (Chameleon, Coherent) with a tunable wavelength served as the excitation source. Emitted fluorescence was separated from the excitation beam using a primary dichroic (Semrock FF705-Di01-49×70) and split using secondary and tertiary long-pass dichroic mirrors (Supplementary Table 1). Emission was detected on photomultiplier tubes through bandpass filters (center wavelength/bandwidth) to image TdTomato or RFP (595/50 nm); FITC (517/65 nm); GFP or Rhodamine-6G (525/70 nm); Qtracker-655 (641/75 nm) and Texas-Red (645/65 nm). Hoechst 33342 and Qtracker-705 emission was detected on photomultiplier tubes without bandpass filters, using the relevant tertiary dichroic to define the collected emission (<488 nm and 560-801 nm). *Z*-stack images were collected from approximately 20 µm below the surface, to a depth of 100 µm (80 µm total imaged depth) moving at 2 µm steps in the z-plane. 100 frames (3.3 s) per plane in z were collected at a scan speed of 30 frames/s. Laser power was increased exponentially as a function of depth. A minimum of four non-overlapping regions were imaged per mouse.

The fluorescence excitation and emission wavelengths for each fluorophore are listed in Supplementary Table 1. All dyes were dosed at 50-100 µL via a retro-orbital injection, with a maximum of 100 µl per eye. Plasma was labeled in Catchup^IVM-red^ mice using fluorescein isothiocyanate (FITC) conjugated dextran (50-100 µL retro-orbitally at the start of imaging 0.02% in saline, 70,000 kDa molecular weight, Thermo Fisher Scientific) To fluorescently label the plasma in Cx3Cr1^+/GFP^-CCR2^+/RFP^ mice, Qtracker 655 or 705 (100 µL retro-orbitally at the start of imaging 0.5 µM in saline, Thermo Fisher Scientific) For strains without endogenous fluorescence (ApoE^-/-^), a combination of dyes was used to label nuclei (Hoechst 33342, 5 mg/mL in saline), various cells and platelets (rhodamine-6G, 0.1% in saline, Thermo Fisher Scientific), and plasma (Texas Red conjugated dextran, 3% in saline, 70,000 kDa molecular weight Thermo Fisher Scientific). Leukocytes were identified by labeling with both Hoechst and rhodamine-6G within the vasculature; platelets were identified by intra-vascular labeling with rhodamine-6G; other cell types (cardiomyocytes, endothelial cells) were identified by their spatial distribution.

**Supplementary Table 1.** Imaging laser and filter wavelengths.

| Mouse strain | Excitation wavelength | Fluorophore | Band-Pass Filter | Secondary dichroic | Tertiary dichroic | Tertiary dichroic |
| --- | --- | --- | --- | --- | --- | --- |
| C57Bl/6 or ApoE<sup>-/-</sup> | 810 nm | Hoechst 33342 | None | 593 | 488 | 801 |
|  |  | Rhodamine-6G | 525/70 |  |  |  |
|  |  | Texas-Red (3%) | 645/65 |  |  |  |
| Catchup-IVM (Ly6G-Cre-TdTom:R26TdTom) | 950 nm | FITC (0.02%) | 517/65 | 560 | 488 | 801 |
|  |  | TdTomato | 595/50 |  |  |  |
| Cx3Cr1<sup>GFP/+</sup> CCR2<sup>RFP/+</sup> | 950 nm | GFP | 525/70 | 605 | 560 | 801 |
|  |  | RFP | 595/50 |  |  |  |
|  |  | QTracker-655 | 641/75 |  |  |  |
|  |  | QTracker-705 | None |  |  |  |

### Quantification of leukocyte dynamics

Individual leukocytes within capillaries were manually identified in two photon image stacks (∼260-s long) using ImageJ. The duration for which each cell was present in a capillary anywhere in the field of view was counted. Leukocytes within larger vessels were not included in the analysis. Accumulated time in region was calculated for each stack as the sum of the duration in view across all identified cells. In some cases, image stacks were split in half and counted independently to allow analysis on memory-limited systems. When split stacks were compared with non-split stacks, all cells in non-split stacks with dwell times spanning the mid-point of the stack were counted as two separate cells to align with the split stack sampling. To prepare representative images in Figure 3, a median filter of 1 pixel was applied, and contrast was adjusted for display using only linear scaling.

### Echocardiography

Transthoracic echocardiography was performed on a VisualSonics Vevo 2100 ultrasound using the MS400 transducer (Visual Sonics Inc) in anesthetized (isoflurane 1-2% in 100% oxygen) mice. Measures of systolic function (ejection fraction, fractional shortening) were made using M-mode images were acquired in short axis at the level of the papillary muscle. The apical 4-chamber view was used to obtain pulsed-wave and tissue Doppler measurements at the level of mitral valve. These were used for measures of diastolic function (E wave, e’ wave, E/e’). All measurements were made using Vevo LAB ultrasound analysis software.

### Administration of antibodies against Ly6G

Antibodies against Ly6G (clone 1A8; 127602, BioLegend) or isotype control (anti-IgG2a kappa, clone RTK2758; 400502, BioLegend) were administered either chronically for the last 4 weeks of heart failure induction (2mg/kg intraperitoneal every 3 days) or once (4mg/kg intraperitoneal) at 15 weeks.

### Cryoinfarction

Anesthesia was induced with ketamine/xylazine (100mg/kg, Ketaset; 10mg/kg, Covetrus, as a combined solution in saline) intraperitoneally and 1-3% isoflurane in 100% oxygen. The heart accessed in the same manner described above for intravital imaging. A lesion on the left ventricular wall was created using targeted focal freezing to mimic tissue damage that occurs in myocardial infarct. A 2mm-diameter copper probe was placed in liquid nitrogen for one minute until fully cooled. The tip of the probe was placed in contact with the heart surface for 5-10 seconds. This process was repeated twice. The tip of a 22G catheter was gently placed in the pleural cavity Ribs were closed by encircling with 6-0 Monocryl. The muscle layers and skin were apposed surrounding the catheter with 6-0 Monocryl. Air was gently withdrawn manually using a 1mL syringe attached to the catheter to reestablish negative pressure in the thorax, and the catheter was removed. Lung pressure measured by the ventilator was monitored as the concentration of isoflurane was gradually reduced. Following the resumption of spontaneous respiration, the endotracheal tube was removed. Post-operatively, animals received buprenorphine 0.1mg/kg every 8 hours (Par Pharmaceuticals) heat support, and wet food for three days.

### Hypoxyprobe assay

Mice were injected with 120mg/kg pimonidazole HCl (Hypoxyprobe Inc, HP-100mg, diluted in saline to 100mg/mL) intraperitoneally 90 minutes prior to deep anesthesia with 4% isoflurane in 100% oxygen. Mice were transcardially perfused with 1X phosphate buffered saline (PBS Sigma Aldrich, item #3813), followed by 4% paraformaldehyde (Thermo Fisher, item #O4042) in PBS. Hearts were removed and stored in 4% PFA overnight, sucrose 30% in PBS for 24-48 hours, cryosectioned in 7 µm slices, and stored at -80°C. Sections were washed with 1% sodium dodecyl sulfate (SDS) in PBS for 5 minutes, and blocking buffer (3% bovine serum albumin; 10% normal goat serum; 0.1% Triton-X100 in PBS) was applied for 30 minutes at room temperature. Sections were overlaid with anti-pimonidazole (Hypoxyprobe Inc, item Pab2627) at a dilution of 1:100 in the same blocking buffer and incubated overnight at 4°C in a humidified chamber. Slides were then washed three times in PBS for 3 minutes each, and overlaid with Alexa Fluor 488 Goat Anti-Rabbit IgG (Thermofisher A-11008) at a concentration of 1:200 in PBS for one hour at room temperature. Slides were washed three times with PBS for 3 minutes each, coverslipped with Vectashield anti-fade mounting medium with DAPI (Vector Labs, item H-1200-10), sealed with clear nail polish (Electron Microscopy Sciences, item #72180), and stored at 4°C until imaging. Images were obtained using a Zeiss LSM 710 confocal fluorescence microscope. Two cohorts of mice were used in this analysis. The sections were stained and imaged at separate dates. The cohorts were normalized by scaling intensities the by mean Hypoxyprobe intensity of all HFpEF samples treated with isotype control antibodies (23.52 for cohort 1 vs. 17.42 for cohort 2).

### Immunohistology

Mice were euthanized via pentobarbital overdose (15mg/mouse via intraperitoneal injection). Prior to cessation of cardiac contractions, mice were transcardially perfused with followed by 4% PFA in PBS. Hearts were removed, sectioned transversely with a razorblade in approximately two equal sections, and stored in 4% PFA. After 24 hours, hearts were transferred to 30% sucrose in PBS for 24-48 hours until they no longer floated in the solution. The apical half of hearts were then embedded in Tissue Tek O.C.T Compound (Electron Microscopy Sciences, item 62550-12), and stored at -80°C until cryosectioning Hearts were cryosectioned in 7-µm thick transverse slices and stored at -80°C until staining. At the time of staining, slides were warmed to room temperature and washed three times in 0.1% Tween-20 in PBS for three minutes each, then washed once with 1% SDS in PBS for 5 minutes. Blocking buffer (3% bovine serum albumin; 10% goat serum; 0.1% Triton-X100 in PBS) was applied for 30 minutes at room temperature. For wheat germ agglutinin and isolectin I4 staining, WGA-Alexa-4 IS88 (1:200, ThermoFisher, item #W11261) and IB4 DyLight 594 (1:100, Vector Laboratories, item# DL-1207) were diluted in Hanks basic salt solution (HBSS, Thermo/Gibco 14025) with calcium chloride (1.26mM or 140g/L). Slides were incubated with this solution for 30 minutes at room temperature, protected from light. Slides were washed once in HBSS with calcium chloride for 1 minute, then three times with 0.1% Tween-20 in PBS for three minutes each. Stained slides were coverslipped with Vectashield anti-fade mounting medium with DAPI (Vector Labs, item H-1200-10), sealed with clear nail polish (Electron Microscopy Sciences, item #72180), and stored at 4°C until imaging. Images were obtained using a Zeiss LSM 710 confocal fluorescence microscope.

### Masson’s trichrome staining

Mice were euthanized and perfused as described above. Tissues were then paraffin embedded, sliced in 7-µm thick sections and stained with Masson’s trichrome by the Cornell University Animal Health Diagnostic Center as previously described^53^.

### Statistical analysis

Statistical analysis was performed using GraphPad Prism for macOS Version 10.2.0 (www.graphpad.com). We used confidence interval of 95% and two-tailed tests with significance set at p < 0.05 Bar graphs show mean and standard deviation as error bars.

### Animal measurements

Mice were weighed weekly using a 0.1 gram scale for the duration of the heart failure induction protocol (15 weeks starting at ages 8-12 weeks). Blood pressure was recorded in lightly anesthetized (1% isoflurane in oxygen) mice using a tail cuff monitoring system (CODA, Kent Scientific). For heart weight/tibial Length (HW/TL) following transcardiac perfusion, hearts were dissected from surrounding tissue. Liquid was drained by carefully squeezing the hearts on tissue paper. Hindlimbs were removed, and tibia extracted bilaterally. After removing excess tissue, the length of the tibias was measured, and recorded as the average of the two measurements. Heart weight (HW) and tibial length (TL) were measured, and the HW/TL was calculated as the ratio of weight (g) to mean tibial length (mm).

### Flow cytometry

Mice were anesthetized with ketamine/xylazine (100mg/kg, Ketaset; 10mg/kg, Covetrus, as a combined solution in saline) intraperitoneally. The heart was accessed between the ribs, and blood was collected from the right ventricle using a 1mL syringe and 27-gauge needle. Blood was immediately transferred to MiniCollect K3EDTA tubes (Greiner Bio-One 450530), which were gently inverted to mix. Mice were subsequently euthanized by pentobarbital overdose. A 100 μL aliquot of whole blood was then added to 2 mL of ammonium-chloride-potassium lysing buffer and gently vortexed. Samples were incubated for 10 minutes at room temperature, protected from light, and then centrifuged at 500 x g for 5 minutes. 2mL of fluorescence-activated cell sorting (FACS) buffer was added to the supernatant. The mixture was gently vortexed and centrifuged again at 500 × g for 5 minutes. The supernatant was removed and combined with 0.5 mL of 2% paraformaldehyde (PFA). Samples were incubated on ice for 15 minutes. An additional 2 mL of FACS buffer was added, and the solution was vortexed and centrifuged at 500 × g for 5 minutes; this wash step was repeated twice more. After the final wash, 0.5 mL of buffer was added to the pellet, and samples were kept on ice until being distributed into a 96-well plate for analysis. Analysis was performed using FloJo software. A granulocyte gate was manually drawn. These cells were then gated for expression of tdTom.

## REFERENCES

1 Jones, J. S., Small, D. M. & Nishimura, N. In Vivo Calcium Imaging of Cardiomyocytes in the Beating Mouse Heart With Multiphoton Microscopy. Front Physiol 9, 969, doi:10.3389/fphys.2018.00969 (2018).

2 Borlaug, B. A. The pathophysiology of heart failure with preserved ejection fraction. Nat Rev Cardiol 11, 507–515, doi:10.1038/nrcardio.2014.83 (2014).

3 Massie, B. M. et al. Irbesartan in patients with heart failure and preserved ejection fraction. N Engl J Med 359, 2456–2467, doi:10.1056/NEJMoa0805450 (2008).

4 Yusuf, S. et al. Effects of candesartan in patients with chronic heart failure and preserved left-ventricular ejection fraction: the CHARM-Preserved Trial. Lancet 362, 777–781, doi:10.1016/S0140-6736(03)14285-7 (2003).

5 Anker, S. D. et al. Empagliflozin in Heart Failure with a Preserved Ejection Fraction. The New England journal of medicine 385, 1451–1461, doi:10.1056/NEJMoa2107038 (2021).

6 Kato, S. et al. Impairment of Coronary Flow Reserve Evaluated by Phase Contrast Cine-Magnetic Resonance Imaging in Patients With Heart Failure With Preserved Ejection Fraction. J Am Heart Assoc 5, doi:10.1161/JAHA.115.002649 (2016).

7 Srivaratharajah, K. et al. Reduced Myocardial Flow in Heart Failure Patients With Preserved Ejection Fraction. Circ Heart Fail 9, doi:10.1161/CIRCHEARTFAILURE.115.002562 (2016).

8 Shah, S. J. et al. Prevalence and correlates of coronary microvascular dysfunction in heart failure with preserved ejection fraction: PROMIS-HFpEF. European heart journal 39, 3439–3450, doi:10.1093/eurheartj/ehy531 (2018).

9 Dryer, K. et al. Coronary microvascular dysfunction in patients with heart failure with preserved ejection fraction. American journal of physiology. Heart and circulatory physiology 314, H1033–H1042, doi:10.1152/ajpheart.00680.2017 (2018).

10 Obokata, M. et al. Myocardial Injury and Cardiac Reserve in Patients With Heart Failure and Preserved Ejection Fraction. J Am Coll Cardiol 72, 29–40, doi:10.1016/j.jacc.2018.04.039 (2018).

11 Mohammed, S. F. et al. Coronary microvascular rarefaction and myocardial fibrosis in heart failure with preserved ejection fraction. Circulation 131, 550–559, doi:10.1161/CIRCULATIONAHA.114.009625 (2015).

12 Schiattarella, G. G. et al. Nitrosative stress drives heart failure with preserved ejection fraction. Nature 568, 351–356, doi:10.1038/s41586-019-1100-z (2019).

13 Zeng, H. & Chen, J. X. Microvascular Rarefaction and Heart Failure With Preserved Ejection Fraction. Front Cardiovasc Med 6, 15, doi:10.3389/fcvm.2019.00015 (2019).

14 AbouEzzeddine, O. F. et al. Myocardial Energetics in Heart Failure With Preserved Ejection Fraction. Circ Heart Fail 12, e006240, doi:10.1161/CIRCHEARTFAILURE.119.006240 (2019).

15 Lyle, M. A., Alabdaljabar, M. S., Han, Y. S. & Brozovich, F. V. The vasculature in HFpEF vs HFrEF: differences in contractile protein expression produce distinct phenotypes. Heliyon 6, e03129, doi:10.1016/j.heliyon.2019.e03129 (2020).

16 D’Amario, D. et al. Microvascular Dysfunction in Heart Failure With Preserved Ejection Fraction. Front Physiol 10, 1347, doi:10.3389/fphys.2019.01347 (2019).

17 Franssen, C. et al. Myocardial Microvascular Inflammatory Endothelial Activation in Heart Failure With Preserved Ejection Fraction. JACC Heart Fail 4, 312–324, doi:10.1016/j.jchf.2015.10.007 (2016).

18 Westermann, D. et al. Cardiac inflammation contributes to changes in the extracellular matrix in patients with heart failure and normal ejection fraction. Circ Heart Fail 4, 44–52, doi:10.1161/CIRCHEARTFAILURE.109.931451 (2011).

19 Paulus, W. J. & Tschope, C. A novel paradigm for heart failure with preserved ejection fraction: comorbidities drive myocardial dysfunction and remodeling through coronary microvascular endothelial inflammation. J Am Coll Cardiol 62, 263–271, doi:10.1016/j.jacc.2013.02.092 (2013).

20 Hulsmans, M. et al. Cardiac macrophages promote diastolic dysfunction. J Exp Med 215, 423–440, doi:10.1084/jem.20171274 (2018).

21 Wang, Y. et al. Wnt5a-Mediated Neutrophil Recruitment Has an Obligatory Role in Pressure Overload-Induced Cardiac Dysfunction. Circulation 140, 487–499, doi:10.1161/CIRCULATIONAHA.118.038820 (2019).

22 Devi, S. et al. Multiphoton imaging reveals a new leukocyte recruitment paradigm in the glomerulus. Nat Med 19, 107–112, doi:10.1038/nm.3024 (2013).

23 Ahn, S. J., Anrather, J., Nishimura, N. & Schaffer, C. B. Diverse Inflammatory Response After Cerebral Microbleeds Includes Coordinated Microglial Migration and Proliferation. Stroke; a journal of cerebral circulation 49, 1719–1726, doi:10.1161/STROKEAHA.117.020461 (2018).

24 McDonald, B. et al. Intravascular danger signals guide neutrophils to sites of sterile inflammation. Science 330, 362–366, doi:10.1126/science.1195491 (2010).

25 Cruz Hernandez, J. C. et al. Neutrophil adhesion in brain capillaries reduces cortical blood flow and impairs memory function in Alzheimer’s disease mouse models. Nat Neurosci 22, 413–420, doi:10.1038/s41593-018-0329-4 (2019).

26 Neupane, A. S. & Kubes, P. Imaging reveals novel innate immune responses in lung, liver, and beyond. Immunol Rev 306, 244–257, doi:10.1111/imr.13040 (2022).

27 Allan-Rahill, N. H., Lamont, M. R. E., Chilian, W. M., Nishimura, N. & Small, D. M. Intravital Microscopy of the Beating Murine Heart to Understand Cardiac Leukocyte Dynamics. Front Immunol 11, 92, doi:10.3389/fimmu.2020.00092 (2020).

28 Lee, S. et al. Real-time in vivo imaging of the beating mouse heart at microscopic resolution. Nature communications 3, 1054, doi:10.1038/ncomms2060 (2012).

29 Kavanagh, D. P. J., Lokman, A. B., Neag, G., Colley, A. & Kalia, N. Imaging the injured beating heart intravitally and the vasculoprotection afforded by haematopoietic stem cells. Cardiovascular research 115, 1918–1932, doi:10.1093/cvr/cvz118 (2019).

30 Jung, S. et al. Analysis of fractalkine receptor CX(3)CR1 function by targeted deletion and green fluorescent protein reporter gene insertion. Molecular and cellular biology 20, 4106–4114 (2000).

31 Saederup, N. et al. Selective chemokine receptor usage by central nervous system myeloid cells in CCR2-red fluorescent protein knock-in mice. PloS one 5, e13693, doi:10.1371/journal.pone.0013693 (2010).

32 Auffray, C. et al. Monitoring of blood vessels and tissues by a population of monocytes with patrolling behavior. Science 317, 666–670, doi:10.1126/science.1142883 (2007).

33 Bajpai, G. et al. Tissue Resident CCR2- and CCR2+ Cardiac Macrophages Differentially Orchestrate Monocyte Recruitment and Fate Specification Following Myocardial Injury. Circulation research 124, 263–278, doi:10.1161/CIRCRESAHA.118.314028 (2019).

34 Jung, K. et al. Endoscopic time-lapse imaging of immune cells in infarcted mouse hearts. Circulation research 112, 891–899, doi:10.1161/CIRCRESAHA.111.300484 (2013).

35 Li, W. et al. Intravital 2-photon imaging of leukocyte trafficking in beating heart. The Journal of clinical investigation 122, 2499–2508, doi:10.1172/JCI62970 (2012).

36 Matsuura, R. et al. Intravital imaging with two-photon microscopy reveals cellular dynamics in the ischeamia-reperfused rat heart. Scientific reports 8, doi:ARTN 15991 10.1038/s41598-018-34295-w (2018).

37 Erdener, S. E. et al. Dynamic capillary stalls in reperfused ischemic penumbra contribute to injury: A hyperacute role for neutrophils in persistent traffic jams. Journal of cerebral blood flow and metabolism: official journal of the International Society of Cerebral Blood Flow and Metabolism 41, 236–252, doi:10.1177/0271678X20914179 (2021).

38 Sequeira, V. et al. Synergistic role of ADP and Ca(2+) in diastolic myocardial stiffness. J Physiol 593, 3899–3916, doi:10.1113/JP270354 (2015).

39 Cruz Hernandez, J. C. et al. Neutrophil adhesion in brain capillaries reduces cortical blood flow and impairs memory function in Alzheimer’s disease mouse models. Nature neuroscience, doi:10.1038/s41593-018-0329-4 (2019).

40 Ali, M. et al. Inhibition of peripheral VEGF signaling rapidly reduces leucocyte obstructions in brain capillaries and increases cortical blood flow in an Alzheimer’s disease mouse model. bioRxiv, 2021.2003.2005.433976, doi:10.1101/2021.03.05.433976 (2021).

41 Bracko, O. et al. Increasing cerebral blood flow improves cognition into late stages in Alzheimer’s disease mice. Journal of cerebral blood flow and metabolism: official journal of the International Society of Cerebral Blood Flow and Metabolism 40, 1441–1452, doi:10.1177/0271678X19873658 (2020).

42 van Heerebeek, L. et al. Diastolic stiffness of the failing diabetic heart: importance of fibrosis, advanced glycation end products, and myocyte resting tension. Circulation 117, 43–51, doi:10.1161/CIRCULATIONAHA.107.728550 (2008).

43 Reeson, P., Choi, K. & Brown, C. E. VEGF signaling regulates the fate of obstructed capillaries in mouse cortex. Elife 7, doi:10.7554/eLife.33670 (2018).

44 Camici, P. G., Tschope, C., Di Carli, M. F., Rimoldi, O. & Van Linthout, S. Coronary microvascular dysfunction in hypertrophy and heart failure. Cardiovascular research 116, 806–816, doi:10.1093/cvr/cvaa023 (2020).

45 Zhu, L. & He, P. fMLP-stimulated release of reactive oxygen species from adherent leukocytes increases microvessel permeability. American journal of physiology. Heart and circulatory physiology 290, H365–372, doi:10.1152/ajpheart.00812.2005 (2006).

46 Boralkar, K. A. et al. Value of Neutrophil to Lymphocyte Ratio and Its Trajectory in Patients Hospitalized With Acute Heart Failure and Preserved Ejection Fraction. The American journal of cardiology 125, 229–235, doi:10.1016/j.amjcard.2019.10.020 (2020).

47 Hage, C. et al. Myeloperoxidase and related biomarkers are suggestive footprints of endothelial microvascular inflammation in HFpEF patients. ESC Heart Fail 7, 1534–1546, doi:10.1002/ehf2.12700 (2020).

48 Kain, V. & Halade, G. V. Role of neutrophils in ischemic heart failure. Pharmacol Ther 205, 107424, doi:10.1016/j.pharmthera.2019.107424 (2020).

49 Kolaczkowska, E. & Kubes, P. Neutrophil recruitment and function in health and inflammation. Nat Rev Immunol 13, 159–175, doi:10.1038/nri3399 (2013).

50 Zhang, X. L. et al. HMGB1-Promoted Neutrophil Extracellular Traps Contribute to Cardiac Diastolic Dysfunction in Mice. J Am Heart Assoc 11, e023800, doi:10.1161/JAHA.121.023800 (2022).

51 Iliakis, C. S. & Wack, A. Never trust a single myeloid marker: Ly6G on repair-promoting lung macrophages. Sci Immunol 9, eadq7306, doi:10.1126/sciimmunol.adq7306 (2024).

52 Piedrahita, J. A., Zhang, S. H., Hagaman, J. R., Oliver, P. M. & Maeda, N. Generation of mice carrying a mutant apolipoprotein E gene inactivated by gene targeting in embryonic stem cells. Proceedings of the National Academy of Sciences of the United States of America 89, 4471–4475, doi:10.1073/pnas.89.10.4471 (1992).

53 Jager, M. C., Choi, E., Tomlinson, J. E. & Van de Walle, G. Naturally acquired equine parvovirus-hepatitis is associated with a wide range of hepatic lesions in horses. Vet Pathol 61, 442–452, doi:10.1177/03009858231214024 (2024).

